# Exon-intron architecture determines mRNA stability by dictating m6A deposition

**DOI:** 10.1101/2022.06.29.498130

**Authors:** Anna Uzonyi, Boris Slobodin, Schraga Schwartz

## Abstract

N6-methyladenosine (m6A), a widespread destabilizing mark on mRNA, is non-uniformly distributed across the transcriptome, yet the basis for its selective deposition is unknown. Here, we uncover that m6A deposition is not selective. Instead, m6A distribution is exclusion-based: m6A-consensus harboring sites are methylated by default, *unless* they are within a window of up to ∼200 nt from an exon-intron junction. A simple model, relying exclusively on presence of m6A motifs and exon-intron architecture allows high accuracy recapitulation of experimentally-measured m6A profiles and of all m6A hallmarks. We further establish that m6A serves as the long-sought mechanism underlying the strong association between exon-intron architecture and mRNA stability. Our findings establish a mechanism by which the memory of nuclear RNA splicing is covalently etched on an mRNA, in the form of m6A, and determines its cytoplasmic stability, with broad implications on the regulation, function, and evolution of both m6A and mRNA stability.

## Introduction

N6-methyladenosine (m6A) is the most widespread modification on mRNA, present at roughly 0.2-0.4% of all adenosines (Perry et al., 1975). A key determinant guiding the RNA methylation machinery towards its targets (the ‘m6A code’) is sequence. m6A is installed at a consensus motif, whose core is typically represented as a DRACH motif (D=A/G/T/ R=A/G, H=A/C/T), which also extends into adjacent nucleotides (Garcia-Campos et al., 2019; Schwartz et al., 2013). Yet, a major feature of m6A distribution, which is not accounted for by sequence, is the strong positional enrichment of m6A within genes. m6A was originally described by us and others to be highly enriched within atypically long internal exons and near stop codons (Dominissini et al., 2012; Meyer et al., 2012). A subsequent study suggested that rather than being associated with the stop codon, m6A was associated with the last intron-exon junction within a gene (which is typically, but not invariably, found in close proximity to the stop codon) (Ke et al., 2015). This prompted multiple studies to explore a potential impact of methylation on splicing, whereby only weak effects were typically observed (Lence et al., 2016; Wei et al., 2021; Xiao et al., 2016). In parallel, alternative potential mechanisms underlying the biased distribution of m6A have been proposed, invoking chromatin marks biased towards the ends of genes and potential involvement of the transcription termination machinery (Abakir et al., 2019; Ke et al., 2015; Yang et al., 2019). However, none of the proposed mechanisms account for the enrichment of m6A near stop codons, nor near last exons nor within long internal exons. The highly biased distribution of m6A has thus remained enigmatic, and the mechanism underlying it largely unknown. Given that m6A plays a well-established role in directing cytoplasmic degradation of mRNA (Dierks et al., 2021; Ivanova et al., 2017; Ke et al., 2017; Kontur et al., 2020; Lasman et al.; Wang et al., 2014; Zaccara and Jaffrey, 2020), elucidating the forces shaping it is pivotal for obtaining a complete understanding of the forces governing mRNA stability.

mRNA levels are shaped by production and degradation. Whereas the factors contributing to production have been intensively studied, the factors determining mRNA degradation rates are understood to a lesser extent. Intriguingly, diverse studies have consistently found that the strongest correlate of transcriptome-wide RNA stability is the ‘exon density’ within the coding region, a ratio between the number of exons within the coding region and the coding region length (Agarwal and Kelley; Agarwal and Shendure, 2020; Clark et al., 2012; Sharova et al., 2009; Spies et al., 2013). The contribution of ‘exon density’ dramatically exceeds that of other mechanisms traditionally implicated in mRNA destabilization, such as miRNA binding sites or AU-rich elements (Agarwal and Kelley; Agarwal and Shendure, 2020), and can alone explain nearly 30% of the variability in degradation rates between different genes (Spies et al., 2013). While this observation suggests a strong connection between exon-intron architecture and RNA stability, its mechanistic basis is completely unknown. Exon-intron architecture of genes has also been associated with additional features of genes including RNA processing, export, and translation (Buchman and Berg, 1988; Dwyer et al., 2021; Le Hir et al., 2001, 2003; Luo and Reed, 1999; Matsumoto et al., 1998; Proudfoot et al., 2002; Zuckerman et al., 2020). The underlying mechanisms are poorly understood, whereby a key gap in our understanding is how the history of splicing in the nucleus is encoded to subsequently drive cytoplasmic fate.

Here, we explore the basis for the region-specific distribution of m6A. On the basis of dozens of distinct backbones and thousands of sequence variants, we find that neither specific sequence elements, nor an intact reading frame, nor RNA splicing, nor chromatin marks, nor a defined distance from termination sites are essential for methylation. Instead, we reveal that that m6A selectivity is not inclusion-based, but rather exclusion based: sites harboring an m6A consensus sequence are methylated by default, *unless* they are within a window of up to 200 nt from an exon-intron junction, in which case the probability of methylation declines with proximity to the junction. We demonstrate that a simple model relying exclusively on presence of an m6A motif and the exon-intron architecture of genes is strongly predictive of experimentally-measured m6A profiles. Our findings are suggestive of a deep mechanistic dependency of mRNA methylation on mRNA splicing, and establish a hard-coded mechanism by which the former serves as a covalently-installed ‘memory’ of the latter, subsequently shaping cytoplasmic stability.

## Results

### A massively-parallel reporter assay for elucidating the sequence determinants of m6A deposition

We previously employed correlative analysis to dissect the m6A consensus sequence, revealing an extended consensus sequence in both human and mouse encompassing a 9-nt window surrounding the methylated consensus (Garcia-Campos et al., 2019). Yet, the m6A consensus motif is not enriched near the stop codon **(Fig. S1a)**, a hallmark of m6A distribution. This raised the possibility that a yet unidentified sequence element might be enriched near the stop codon or the last exon and could be driving methylation. To explore this possibility, we sought to establish a massively-parallel reporter assay (MPRA) to interrogate the sequence determinants governing methylation. As a proof of concept, we first selected a 101-bp region from the human SLC25A3 3’ UTR **(Fig. 1a, S1b)** displaying very strong and reproducible enrichment in m6A-seq experiments, and containing a single, highly reproducible m6A site identified by 6 out of 6 m6A-miCLIP experiments (Chen et al., 2019); While this sequence contains a few additional DRACH motifs, no additional m6A sites had been identified by even a single miCLIP experiment within this region. This region was cloned either in its original form (‘WT’) or with a point mutation in the primary m6A consensus sequence (‘mut-main’) into a plasmid backbone as a 3’ UTR element downstream of GFP. Application of m6A-seq2 (Dierks et al., 2021) to poly(A)-selected mRNA extracted from the transfected cells revealed roughly 5.5-fold enrichment (IP/input) of the WT construct, while the ‘mut-main’ counterpart exhibited a substantially reduced enrichment **(Fig. 1a, S1b)**. Given that we still observed residual enrichment in the point-mutated construct, we designed two additional constructs - one in which we abolished all secondary m6A consensus sequences but not the ‘primary’ one (‘mut-secondary’) and one in which we abolished all m6A consensus sequences (‘mut-all’). We found that the ‘mut-secondary’ constructs were enriched at levels similar to the ‘mut-main’, whereas enrichment was entirely abolished in the ‘mut-all’ constructs **(Fig. 1a, S1b)**. These analyses thus establish that m6A peaks are faithfully recapitulated within our plasmid-based reporter system, and suggest that at least in the examined example, a single ‘m6A peak’ is due to multiple, closely adjacent methylation sites.

**Figure 1.**
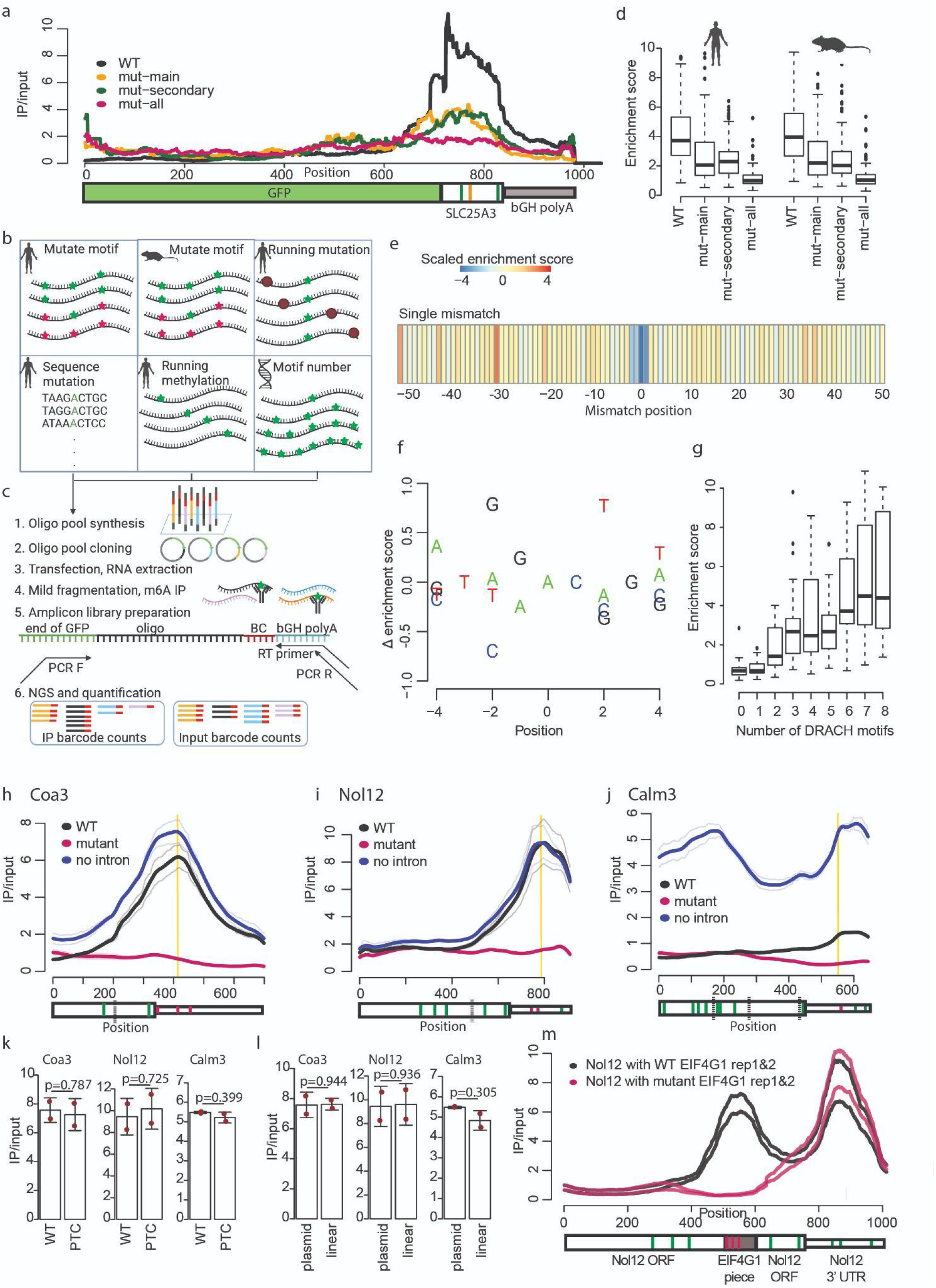
Relative position within a gene, chromatin, reading frames and introns are not required for m6A formation. **a)** IP/input ratios of GFP with a 3’ UTR insert from SLC25A3. The WT construct contains a 101 bp long stretch of SLC25A3, with a GGACT sequence detected in 6 out of 6 miCLIP experiments. The mut-main construct has the main, annotated motif mutated to GGTCT at the position indicated in yellow within the gene model. The mut-secondary construct has the main motif unchanged, but additional nearby DRACH motifs at positions indicated in green are point mutated. The mut-all construct has all DRACH motifs point mutated. To calculate the IP/input ratio, IP reads at each position were normalized to the total number of IP reads mapped to the plasmid. Input reads at each position were normalized to the total number of input reads mapped to the plasmid. Normalized IP reads were divided by normalized input reads. Quantification at the main m6A motif position is shown in Fig. S1b. **b)** Depiction of the library subsets analyzed in Fig. 1 and Fig. S1. **c)** Experimental pipeline of the m6A measurement of the oligo library pool. Oligo pool was planned and synthesized, followed by cloning into a plasmid. RNA was extracted from cells transfected with the library pool and subjected to mild fragmentation. Two rounds of m6A immunoprecipitation was followed by a site-specific reverse transcription and subsequent preparation of amplicon libraries and next generation sequencing. **d)** Enrichment score of WT and control sequences (as in a) on the basis of 200 human and 200 mouse sequences. Outliers with an enrichment score over 10 are not shown. Box plots correspond to the median, Q1 and Q3, whiskers mark Q1-1.5 IQR and Q3+1.5 IQR. **e)** Heatmap of enrichment score z-scores in the running single nucleotide point mutation. For the additional sets of mutations of three or five consecutive bases, see Fig. S1e. **f)** Delta (Δ) enrichment scores of the motif sequence set, on the basis of permutation of the extended 9-mer consensus motif in four human mut-secondary sequences. The methylated A corresponds to position 0. Each value is a mean of all sequences with a certain base at a given position. To calculate the delta enrichment scores at each position, from the mean enrichment score of a given nucleotide we subtracted the mean enrichment score of any other nucleotides. These scores were averaged for all the different motif permutations for each position. **g)** Enrichment of the motif number subset. Box plots correspond to the median, Q1 and Q3, whiskers mark Q1-1.5 IQR and Q3+1.5 IQR. The boxes correspond to sequences containing 0 to 8 m6A consensus motifs. Each box is based on 18 synthetic sequences with different extended 9-mer consensus motifs. **h)** IP/input ratio over the Coa3 coding region and 3’ UTR. Thin lines are measurements from two biological replicates. Thick lines are the average of these measurements with a 10% Loess fitting. In the bottom annotation, the thick box marks the CDS, the thin box marks the 3’ UTR. Black dotted lines mark the exon-intron junctions, green and red stripes mark strong DRACH motifs (GGACC, AGACA, TGACT, AGACT, GAACT, GGACA, GGACT), where the red ones were modified in the mutant constructs and the green ones were left unchanged. The vertical yellow stripe indicates the main DRACH motif and the position analyzed in the right panel. Quantification of the IP/input ratio of the main 3’ m6A site is shown in Fig. S1h. **i)** Measurements for the Nol12 gene. All annotations are according to panel h. **j)** Measurements for the Calm3 gene. All annotations are according to panel j. **k)** IP/input ratios at the ‘main’ site across each of the three constructs in the WT sequences (lacking an intron) in comparison to counterparts into which a premature termination codon (PTC) was introduced. All constructs lack introns. Red dots mark individual measurements, the height of the bars mark the average and the whiskers represent the standard deviation of the mean. P-values for two-tailed Student’s t-test are shown. **l)** IP/input ratios at the ‘main’ site across each of the three constructs in the WT constructs (lacking an intron) which were transfected as plasmid into cells in comparison to corresponding measurements in the same constructs that had been introduced into cells as linear DNA following amplification from the plasmids. All constructs lack introns. Red dots mark individual measurements, the height of the bars mark the average and the whiskers represent the standard deviation of the mean. P-values for two-tailed Student’s t-test are shown. **m)** IP/input coverage of constructs with a 101 nt long EIF4G1 derived piece (WT or DRACH depleted mutant) cloned within the open reading frame (ORF) of the intronless Nol12 gene. Based on two biological replicates. Wide box marks the open reading frame, the thin box marks the 3’ UTR. m6A motifs are marked in green over the Nol12 gene. The DRACH motifs in the EIF4G1 fragment are marked in magenta.

We next designed a series of 7584 different sequences, each 101-bp in length. These sequences included (1) a ‘Mutated motif’ series, comprising 200 human and 200 mouse sequences with a strongly methylated motif, and the corresponding mut-main, mut-secondary and mut-all sequences, similarly to the above-described reporter, (2) a ‘Running mutation’ series on the basis of 10 human mut-secondary sequences, where one, three or five consecutive nucleotides are mutated to their complement, running from the beginning to the end of the sequence, (3) a ‘Sequence mutation’ series of systematic mutations of the extended, 9-mer consensus motif on the basis of four human mut-secondary sequences, (4) a ‘Running methylation’ series on the basis of 10 human mut-secondary sequences, moving a single, DRACH motif from the beginning to the end of the sequence and, (5) a ‘Motif number’ series, consisting of fully synthetic sequences with 0-8 repeats of 18 different 9-mer consensus motifs, consisting of both a DRACH core and an A and a U at positions -4 and +4 respectively (Garcia-Campos et al., 2019) **(Fig. 1b)**, in addition to 4371-sequences that will be described elsewhere. All sequences were cloned as a pool downstream of the sequence of GFP and the resulting plasmid pool was transfected into HEK293T cells **(Fig. 1c)**. RNA extracted from the cells was subjected to Poly(A)-selection, mild fragmentation, and m6A-IP followed by targeted amplification of the insert and library construction. Each sequence was associated with an enrichment score, calculated as the ratio between the number of reads originating from this sequence upon m6A-IP in comparison to the ‘input’ control, normalized to the median enrichment score of all human, mouse, and synthetic constructs lacking an m6A consensus motif in the library. Enrichment scores were highly reproducible internally (Pearson R (R_p_) = 0.95, **Fig. S1c**), as evaluated on the basis of 150 sequences that had been integrated twice into the pool with different barcodes; This analysis also rules out substantial contribution by the barcode associated with each sequence (of note: the barcodes were designed to lack any ‘AC’ motif, so as to prevent methylation). Enrichment scores were also reproducible between biological replicates (R_p_ = 0.82, **Fig. S1d**).

We first analyzed the wild-type human and mouse ‘Mutate Motif’ series. Reassuringly, enrichment scores of methylated sequences were higher than their point-mutated counterparts **(Fig. 1d)**. Similar to our pilot experiments, we found that a series of single point-mutants at the ‘main’ methylation site only gave rise to a partial drop in methylation levels. A similar level of reduction was also achieved in a series in which all secondary methylation consensus sequences - but not the primary one - were point-mutated **(Fig. 1d)**. Complete elimination of m6A was only observed following elimination of both primary and secondary sites **(Fig. 1d)**. These findings thus suggest that methylation at multiple adjacent sites, all of which additively contribute to the amplitudes of a single ‘m6A peak’, is the rule rather than an exception, at least under the surveyed conditions.

We next analyzed the ‘running mutation series’, to explore whether sequences other than the methylation consensus sequence might be required for methylation. This analysis revealed that single nucleotide mutations that disrupt positions -2, -1, 0, or +1 abolished methylation **(Fig. 1e)**. A reduction was also evident in the 3mer and 5mer mutation series, for any window that reached these positions **(Fig. S1e)**. Other than the sequence window immediately surrounding the methylated site, no other position consistently impacted methylation.

To unravel the relative contribution of nucleotide composition at the different positions surrounding the methylation site, we next analyzed the systematic ‘sequence mutation’ series in which the bases surrounding methylation sites were systematically point-mutated. On the basis of these *functional* perturbations we could reconstruct de-novo the extended methylation consensus sequence that we had previously identified merely on the basis of an *associative* analysis **(Fig. 1f)**. This functional reconstruction de-novo uncovered both the requirement for a ‘DRACH’ motif at positions -2 to +2, as well as a preference for an A and a U at positions -4 and +4, respectively (Garcia-Campos et al., 2019).

Finally, we analyzed the ‘running methylation series’, in which the relative position of the methylation site within the sequence was shifted. Of note, this series effectively modulates both the distance of the methylation site with respect to the stop codon and with respect to the transcription termination sites. We observed methylation throughout all constructs in this series, without any obvious positional effects **(Fig. S1f)**.

These analyses thus all suggest that, at least in the surveyed reporters, the determinants *required* for methylation are to a large extent confined to the methylation consensus sequence. To explore whether the consensus sequences might also be *sufficient* for achieving methylation, we analyzed a series of completely synthetic sequences into which we had installed a varying number of methylation consensus motifs. Indeed, we found that these synthetic sequences were progressively and efficiently enriched with an increasing number of methylation consensus sequences **(Fig. 1g)**.

Collectively, the above analyses thus suggest that within the surveyed reporters relying on expression of intronless GFP (1) a methylation consensus sequence is both required and sufficient for achieving methylation, (2) multiple consensus sites contribute additively to measured methylation ‘peaks’, and (3) neither the neighboring positions nor the relative location of a methylation consensus plays a significant role in modulating methylation levels.

### Splicing, reading frames, chromatin and proximity to 3’ end are not required for methylation

If the methylation consensus sequence is the only factor driving methylation, then why is m6A enriched in long exons or near stop codons, or near last exon junctions? One possibility is that methylation consensus motifs are enriched in these regions, but this is evidently not the case (**Fig. S1a**). We reasoned that the answer to these questions might reside within more distal elements within genes which cannot be systematically perturbed at high throughout, and instead require tailored mutations at specific positions within the backbones.

Motivated by previous results that the ‘stop-codon’ associated peaks correlate with the last intron-exon junction (Ke et al., 2015), we first sought to assess whether RNA splicing is required for methylation. We selected three mouse genes with strong peaks in the stop codon vicinity and with a relatively short last intron (to facilitate cloning): Coa3, Nol12, and Calm3. The exons encoding each of the genes, in addition to the last intron (preceding the methylation site) were cloned into plasmids either in their wild-type form (‘WT’), or with point-mutation of the suspected methylation motifs (‘point-mut’). In addition, we cloned versions of the same sequences but without the last introns (‘no-intron’). In one of the 3 cases (‘Calm3’) the introns were sufficiently short to enable us to include three introns of this gene in the WT and ‘point-mut’ form. Application of m6A-seq2 to all 9 constructs, following their transfection into MCF7 cells, revealed that the ‘WT’ constructs recapitulated the ‘stop-codon’ proximal enrichment of the endogenous constructs **(Fig. 1h-j, S1h)**, which was dramatically reduced in the point-mutated counterparts **(Fig. 1h-j, S1h)**. However, no reduction was observed in the intron-less constructs - indeed, increased methylation levels were observed in two out of the three cases, which was statistically significant in the case of Calm3 **(Fig. 1h-j, S1h)**. These results thus indicate that splicing is not required for methylation.

Given the association of the peaks with stop codons, we next sought to assess whether an intact reading frame was required for methylation. Using the three ‘no-intron’ constructs as a basis, we introduced a stop codon early on in the reading frame. However, in none of these constructs did we observe an impact on RNA methylation **(Fig. 1k)**. These experiments thus rule out a model by which methylation is directed via a proximal stop codon.

In recent years multiple studies have linked m6A deposition to chromatin, via diverse mechanisms (Barbieri et al., 2017; Huang et al., 2019; Knuckles et al., 2017; Li et al., 2020; Liu et al., 2020a). In particular, it was suggested that the 3’ biased localization of m6A could be directed by a specific chromatin mark (H3K36me3) displaying a 3’ bias within gene bodies (Huang et al., 2019). To explore whether chromatin was required for directing RNA methylation in the surveyed contexts, we PCR-amplified each of the three genes and transfected the linear chromatin-free PCR products into cells. Methylation profiles in this experiment were essentially identical to the ones observed via transfection of the plasmids **(Fig. 1l)**, ruling out that chromatin is required for 3’ biased m6A formation in the surveyed contexts.

To explore whether the relative location of the methylation site within the gene was required for methylation, we introduced the methylated region or a point-mutated counterpart derived from the 3’ UTR of EIF4G1 into the gene body of the Nol12 no-intron construct, thereby generating a hybrid gene comprising the EIF4G1-derived methylatable region in the gene body and the original Nol12 derived region in the 3’ UTR. Despite the shift into a completely different region, the EIF4G1 site got methylated strongly in WT, but not in the point-mutated counterpart, while the Nol12-derived site was equally methylated in both **(Fig. 1m)**. Thus, presence within a distal region within a gene is not a requirement for methylation.

Our results thus far suggest that methylation does not require any particular sequence other than the methylation motif, nor splicing, nor a reading frame, nor a fixed distance from a polyadenylation site, nor a specific chromatin conformation, nor a relative position within a gene, and that methylation consensus sequences can be sufficient for methylation - rendering the question of the determinants giving rise to the region-specific deposition of m6A all the more enigmatic.

### m6A topologies are driven via exclusion from intron-exon junction proximal regions

In the above experiment we noted that removal of introns led to increased m6A signal **(Fig. 1h-j, S1h)**. This was particularly pronounced in the case of Calm3, a gene harboring multiple introns, but subtle increases were evident in Coa3 and Nol12, in both of which the m6A enrichment increased and also spread over larger regions of the genes, suggesting methylation at additional sites. In the intron-less construct of Calm3 where this was particularly pronounced, substantial enrichment was observed within the CDS, in a region corresponding to the end of exon 1 and beginning of exon 2, which harbored a high density of DRACH sites **(Fig. 1j)**. Such enrichment was not observed in the intron harboring construct, where the methylation signal was confined to the 3’ UTR region (exon 4). The enrichment pattern within the intron-less construct led us to wonder whether it could be de-novo predicted simply on the basis of DRACH sites along it. To explore this, we established a simple model of m6A formation (‘m6Apred-1’) according to which every ‘eligible’ m6A site is methylated at a fixed level. Eligible m6A sites were defined as ones harboring any of the seven most prevalent consensus sequences (GGACC, AGACA, TGACT, AGACT, GAACT, GGACA, GGACT) (Schwartz et al., 2014). To allow a direct comparison between our model and m6A-seq, in which the signal of an m6A site is spread as a ‘peak’ over a window surrounding the methylated site, we modeled each predicted m6A site as a gaussian centered over a 200-bp window surrounding the methylation site. The predicted methylation level of each position was defined as the sum of predicted signal overlapping that position (**Fig. 2a-top**). Remarkably, the m6Apred-1 predicted methylation profile in Calm3 displayed an excellent correlation with measurements in the intron-less context (Spearman R_s_=0.89, P<2.2e-16) (**Fig. 2a-bottom**), but no correlation with the corresponding measurements in the intron-harboring context (R_s_=-0.06, P=0.1), in which the measured signal was confined only to the end. Of note, the in-silico reconstituted methylation profile in Calm3 is based on 11 different eligible DRACH motifs distributed along the length of this gene, all of which are predicted to undergo methylation by this model.

**Figure 2.**
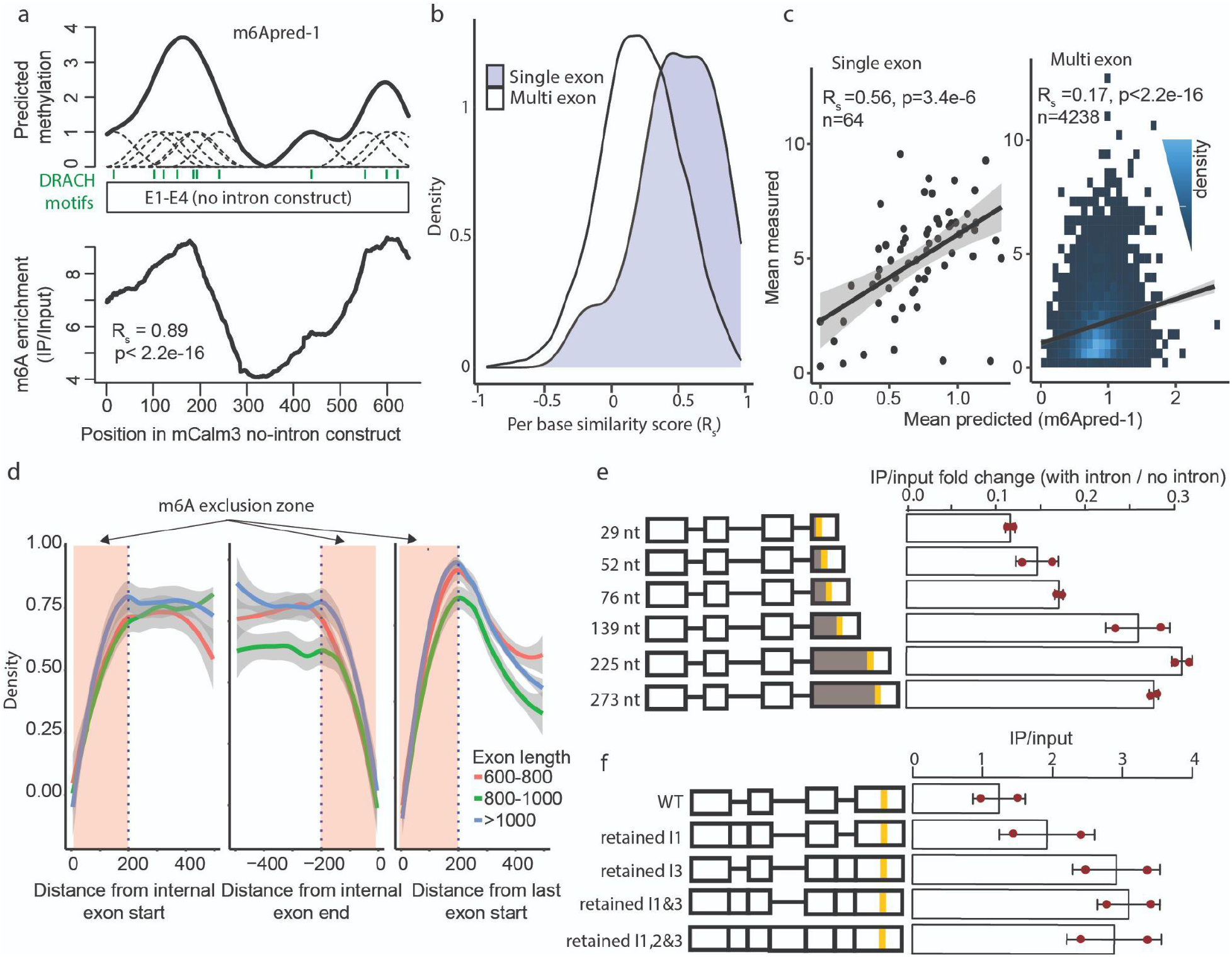
Proximity to intron-exon junctions inhibits m6A formation. **a)** (top) Depiction of m6Apred-1 predictions on the basis of the Calm3 gene. Every eligible DRACH motif (depicted as green bars) is modeled as a Gaussian, whereby the sum of all Gaussians overlapping a position defines its predicted methylation level. (bottom) Measured IP/input values along the Calm3 gene are shown. The indicated correlation reflects the per-base correlation between the predicted (top) and measured (bottom) levels of methylation along the Calm3 gene. **b)** Distribution of per-base similarity scores within single-exon and multi-exon genes. **c)** Correlation between the mean predicted levels of methylation (mean m6Apred-1 values along gene bodies) and corresponding measurements (IP/Input in A549 cells), shown for both single-exon and multi-exon genes. **d)** Left panel: density of detected m6A sites by the distance from the start of internal exons. Middle panel: density of detected m6A sites by the distance from the end of internal exons. Right panel: density of detected m6A sites by the distance from the start of last exons. Blue dashed line marks 200 bp distance from the nearest exon-intron junction. Data is based on an assembly of human miCLIP datasets (Chen et al., 2019). **e)** Left panel: schematic of constructs with different distances between the last splice junction and the strong 3’ m6A motif, on the basis of the Calm3 gene. Boxes mark exons, horizontal lines mark introns. Vertical yellow lines mark the position of the quantified DRACH motif. The distance between the quantified 3’ UTR DRACH motif and the last splice junction is marked in grey. Right panel: Bar plot comparing the methylation levels of the 3’ UTR DRACH site of the WT Calm3 construct, and corresponding constructs with reduced or increased distance between the last splice junction and the quantified site. For the quantification of input and IP signal, only reads beginning at most 10 bp upstream of the modification site were used, to avoid contribution of signal from a more upstream consensus motif (see Methods). Quantification shows the ratio of enrichment between the intron harboring and intronless constructs. The separate (with intron, without intron) quantifications can be found in Fig. S2e. Red dots mark individual measurements, the height of the bars mark the average and the whiskers represent the standard deviation of the mean. **f)** Left panel: schematic of constructs retaining introns 1, 3, 1&3 or all, on the basis of the Calm3 gene. All graphical parameters are according to panel e. Right panel: Bar plots comparing the methylation levels 3’ UTR DRACH site of the WT Calm3 construct, as well as corresponding constructs with retention of intron 1, 3, 1&3 or 1,2&3. Red dots mark individual measurements, the height of the bars mark the average and the whiskers represent the standard deviation of the mean. For the quantification, 5’ read starts up to 40 nt upstream were summed.

To explore whether DRACH-motif distribution along genes was sufficient for predicting methylation patterns in intron-less genes transcriptome-wide, we applied m6Apred-1 to all ∼20,000 canonical UCSC human gene models. We then employed two metrics to assess the ability of m6Apred-1 to capture the variability in experimentally-derived m6A signal in human A549 cells (Schwartz et al., 2014) both *within* genes and *between* genes. For assessing the variability within genes, we employed the per-base Spearman correlation between predicted and measured (IP/input) profiles, as above. To explore variability between genes, we assessed the correlation between mean predicted levels per gene and mean measured (IP/input) enrichment levels. Consistently with our above results, we found that both the variability in m6A levels within genes and between genes was well-captured by m6Apred-1 in intron-lacking genes, but not in intron harboring ones. The median per-base agreement between measured and predicted values was 0.5 in intronless genes (n=64), but 0.21 in intron-harboring ones (n=4238) **(Fig. 2b)**. In parallel, there was high agreement between mean m6Apred-1-based profiles and m6A measurements in intron-lacking genes (R_s_=0.56), but poor agreement in intron-harboring genes (R_s_=0.17) **(Fig. 2c)**. These results thus suggest that in the absence of introns, m6A accumulation to a large extent mirrors the distribution of methylation consensus sequences, but that introns have an inhibitive impact on m6A formation.

This set of results prompted us to explore the following model: rather than assuming that methylation is *off* by default and seeking to identify factors giving rise to it, we hypothesized that an opposite model might be true, in which methylation is *on* by default *unless* it is in proximity to an intron-exon junction.

This model predicts that m6A be depleted from regions proximal to exon-intron junctions. To assess whether this is the case, we examined the distribution of 81,518 methylation sites, identified in human cells at single-nucleotide resolution by six m6A-miCLIP experiments (Chen et al., 2019), with respect to the intron-exon junction. Remarkably, we found m6A depleted from both the 5’ and 3’ ends of long internal exons, as well as from the 5’ of last exons. Specifically, we observed a progressive depletion of m6A over a ∼200 bp window from intron-exon (or exon-intron) junctions **(Fig. 2d, S2a)**. For instance, considering internal exons >600 nt, 32 putative m6A sites resided within the 25 bp immediately following the intron-exon junction, whereas 262 putative sites resided in an equally sized window 269 bp from the junction. The size of this ‘m6A exclusion zone’ was fixed, and did not depend on the length of the internal exon **(Fig. 2d, S2a)**. We furthermore repeated this analysis in mouse, on the basis of 25,879 m6A sites identified via miCLIP (Linder et al., 2015), and made similar observations **(Fig. S2b)**. We also observed the same trend on the basis of a recently published bulk DART-seq (Tegowski et al., 2022) dataset **(Fig. S2c)**, where RNA methylation is detected via an antibody independent manner, excluding that these results could be due to potential biases of antibody-based enrichment. These results thus suggest the relative exclusion of methylation sites from nucleotides residing up to 200 nt from a splice junction. These findings are consistent with previous findings which also noted depletion of m6A from the vicinity of exon-intron junctions (Ke et al., 2017).

To experimentally confirm that distance from the junction causally impacts m6A formation, we next modulated the distance between the Calm3 methylation site and the last intron-exon junction, by either deleting chunks of sequence of varying lengths from the sequence between the splice junction and the m6A site or by introducing additional sequences (lacking DRACH motifs) between the splice junction and the methylation site. Identical modulations of sequence were performed both in the intron-harboring and in the intron-lacking constructs. Consistent with our observations, we found that the distance between the methylation site and the splice junction strongly impacted m6A levels, with increased distances giving rise to increased relative levels, up to a distance of 225 bp **(Fig. 2e, S2e)**. To further explore this, we introduced a 6 bp deletion at the donor site of intron 3, which led to intron 3 retention **(Fig. 2f, S2d)**, giving rise to a 361-bp distance between the methylation site and the nearest junction. Consistent with our expectations, this perturbation gave rise to a strong increase in methylation levels of the 3’ UTR site **(Fig. 2f)**. As a control, we disrupted the splicing of intron 1 leading to its retention, which had a much milder effect. Disruption of splicing of introns 1 and 3, or of all introns, gave rise to a signal similar in magnitude to disruption of only exon 3. This line of experiments also allows to rule out a possibility that methylation is not inhibited by the splicing out of an intron, but instead by proximity to intronic sequence, as in this case the distance to the intronic sequence is maintained and only splicing is inhibited. Thus, these experiments lend strong experimental support to the inhibitory impact of proximity to a splice site on mRNA methylation.

### Transcriptome-wide m6A topologies can be quantitatively reconstituted in-silico on the basis of junction-exclusion based model

Our results thus far strongly suggest that two components are *required* for m6A deposition: a consensus sequence, and a minimal distance from a splice junction. We next sought to assess whether these two parameters are *sufficient* for in-silico transcriptome-wide reconstitution of m6A landscapes. To address this, we developed the ‘m6Apred-2’ model as an expansion of m6Apred-1, incorporating two considerations: (1) Every ‘eligible’ m6A site is methylated (as in ‘m6Apred-1’), (2) unless it is within an ‘m6A exclusion zone’, a region spanning a fixed size (‘ExclusionZoneSize’) from the exon/intron junction (**Fig. 3a-top**). The ‘‘ExclusionZoneSize’ was set at 100 nucleotides, roughly midway within the region depleted of m6A **(Fig. 2d)**. Of note, for intronless genes (but not for multi-exon genes), m6Apred-1 and m6Apred-2 predictions are identical. Encouragingly, when we applied m6Apred-2 to the intron-harboring version of Calm3, the constraint on distance from a junction only allowed 3 DRACH motifs in the (long) terminal exon to undergo methylation, whereas the 8 remaining DRACH motifs residing within the short internal exons were no longer considered eligible. This resulted in a high agreement with experimental measurements, now also in an intron-harboring context (R_s_=0.81, **Fig. 3a-bottom**).

**Figure 3.**
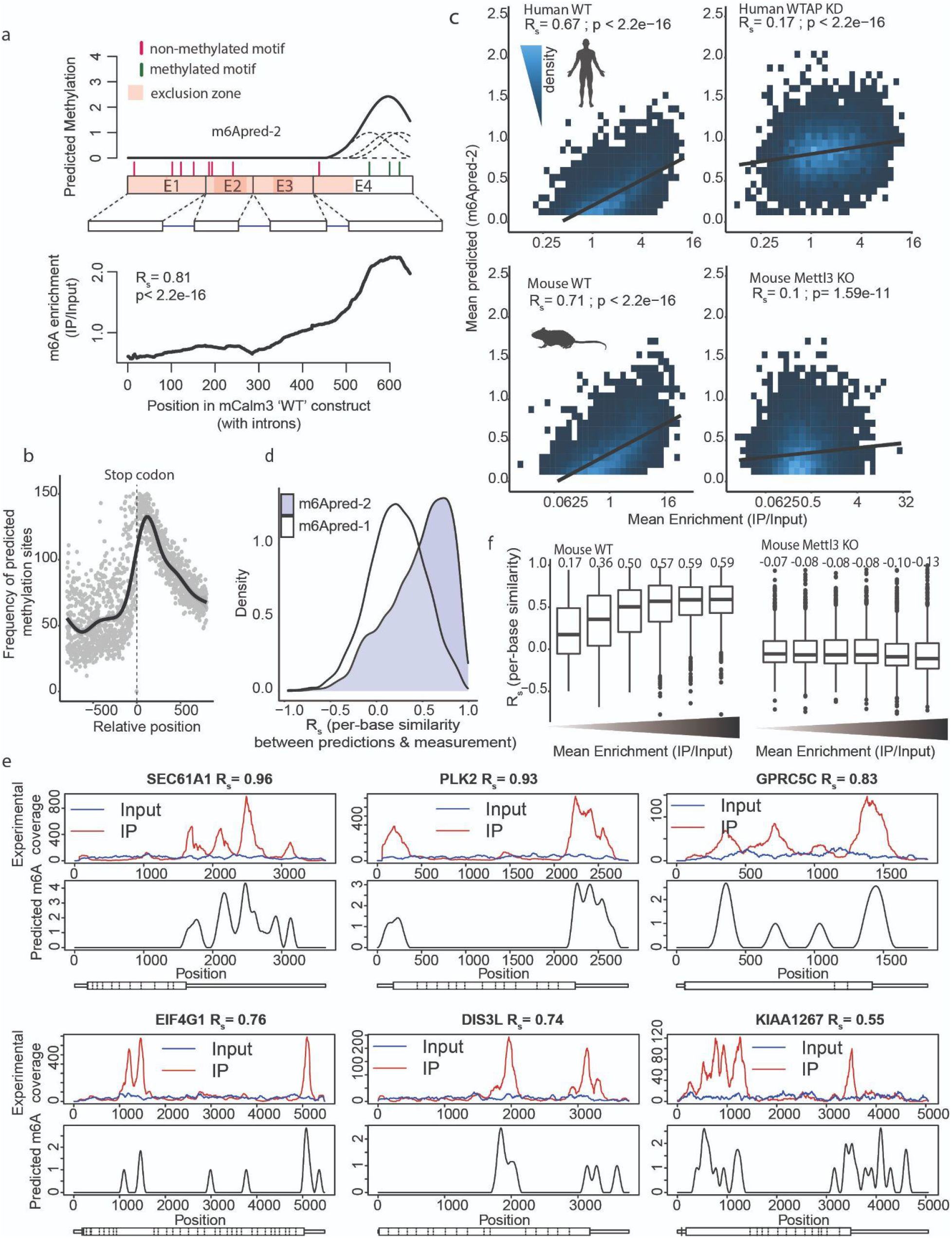
M6A topologies can be de-novo predicted based on motif and exon-intron architecture. **a)** (top) Development of m6Apred-2, relying on eligible DRACH motifs and exon-intron architecture, illustrated on the basis of the Calm3 gene. Eligible m6A sites are depicted as lines, drawn in red if they are within an m6A exclusion zone (within 100 bp of an exon-intron junction, depicted as a transparent box in peach) or green otherwise. Each site is modeled as a gaussian, and the sum of the gaussians at each position define its predicted levels of methylation. (bottom) Measurements of m6A (IP/input) for the ‘WT’ Calm3 construct (harboring agreement). The depicted correlation is the per-base correlation between the measured levels and the predicted ones (top). **b)** Distribution of m6Apred-2 based predicted methylation sites with respect to the stop codon, displaying a strong enrichment around the stop codon, recapitulating to the ‘classical’ m6A topography. Black line shows Loess fitted average. **c)** Agreement between predicted (m6Apred-2) and measured (IP/input) m6A values in human A549 cells (top) and mouse embryonic stem cells (bottom), in both WT cells (left) and cells depleted of methylation components, as indicated (right). Spearman correlations and P values are shown. **d)** Distribution of per-base similarity scores between predicted and measured (A549 cells) m6A levels on the basis of m6Apred-1 (white) and m6Apred-2 (purple). **e)** Predicted and measured m6A levels across six human genes. Top panels display the IP and input signal and bottom panels display the m6Apred-2 based prediction along the gene. The per-base correlation between predicted and measured values is shown at the top of each panel. **f)** Boxplots depicting the distribution of the per-base similarity scores across mouse genes, binned into 6 equally sized groups on the basis of the experimentally measured m6A levels (IP/Input). Distributions are shown for both WT mESC cells (left panel) and for METTL3 KO counterparts (right panel).

Application of this model to 20,102 human gene-models yielded 262,326 predicted methylation sites, with a mean of ∼13 predicted sites per gene. These sites were distributed over 76,599 predicted ‘peaks’ (whereby each peak was defined as a consecutive stretch of bases with a predicted methylation signal >0). An average peak comprised 3.4 methylated sites, and spanned 373 nt, consistent with our results based on the massively-parallel reporter assay that multiple methylated sites per peak were the rule, rather than an exception. We obtained similar results across 20,954 mouse genes, where an average of 12.8 sites per gene distributed across 68,893 peaks.

To evaluate the performance of this model, we assessed its ability to capture m6A features at three levels: m6A ‘meta-features’, variability in m6A levels between genes, and variability in m6A levels within genes. We first found that the predicted m6A profiles recapitulate all previously reported m6A meta-features, including the enrichment of m6A near stop codons **(Fig. 3b)** as well as the relative depletion of m6A from highly expressed genes **(Fig. S3a)**. Next, we explored the ability of the model to predict diversity in methylation *between* genes. We observed a striking correlation (R_s_=0.67) between experimentally measured m6A levels in A549 cells (Schwartz et al., 2014) and mean m6Apred-2 predictions **(Fig. 3c, top left panel)**. Importantly, this correlation coefficient dropped substantially (R_s_=0.17) when predicted values were compared against m6A measurements in A549 cells which had been pre-treated with siRNAs targeting WTAP **(Fig. 3c, top right panel)**, a critical member of the methyltransferase complex whose abolishment substantially reduced methylation levels (Schwartz et al., 2014). We furthermore made essentially identical observations in mouse embryonic stem cells, where we observed excellent agreement between predicted methylation densities and experimentally measured counterparts (R_s_=0.71, **Fig. 3c, bottom left panel**). In this experiment we also had access to m6A measurements in METTL3 knockout cells, devoid of methylation, in which case the correlation with the predicted levels was nearly entirely abolished (R_s_=0.1, **Fig. 3c, bottom right panel**), ruling out that promiscuous binding by the antibody somehow accounts for these associations. Thus, roughly 45-50% of the variability in experimentally-measured m6A levels at the gene levels are captured by our model.

Finally, m6Apred-2 also performed remarkably well in de-novo recapitulating experimentally-measured m6A distribution *within* genes, with genes displaying remarkably high similarity scores between measured and predicted values **(Fig. 3d)**. Roughly 15% of the genes displayed excellent similarity scores (R>0.8) across gene bodies - in such cases, measured m6A profiles are nearly perfectly recapitulated using the simple, above-defined model. Another ∼26% of the cases displayed correlations between 0.6-0.8, and ∼22% displayed correlations ranging from 0.4-0.6. Examples for genes displaying varying similarity scores are displayed in **Fig. 3e** and in **Fig. S4**. Median similarity scores in an intron-containing context were 0.54, marking a dramatic improvement over m6Apred-1 based predictions which achieved a median score of 0.21 **(Fig. 2b)**.

The availability of position-specific predicted methylation profiles allowed us to explore whether there were any properties of genes, or regions within genes, in which our model performed suboptimally. These explorations gave rise to several observations: (1) Genes with lower methylation levels generally showed substantially poorer similarity scores. For instance, the most lowly methylated genes displayed median similarity scores of 0.17, whereas the most highly methylated bin displayed similarity scores of 0.59 **(Fig. 3f, S3b, S4)**. This is anticipated: in lowly-methylated genes the relative levels of signal (true methylation) to background (non-specific binding) is low, and hence in such genes a more significant portion of the m6A quantifications reflect ‘noise’ which is not captured by our model. (2) M6Apred-2 tended to underestimate m6A signal in 5’ UTR regions **(Fig. S3c,d)**, likely due to the fact that a substantial portion of m6A enrichment in the 5’ UTR is not due to m6A, but instead due to a distinct modification - m6Am - which is installed by a different machinery at non-DRAC related sites (Akichika et al., 2019; Keith et al., 1978; Mauer et al., 2016; Schwartz et al., 2014), and hence not accounted for by the model. (3) Conversely, m6Apred-2 tended to overestimate m6A levels in the 3’ UTR **(Fig. S3c,d)**, which is likely, at least partially, due to the difficulty in annotating 3’ UTRs. 3’ UTR annotations often rely on the ‘longest’ observed isoforms, but the actually utilized 3’ UTRs is often substantially shorter, as was also observed in our data **(Fig. S3e)**. In such cases, methylations are predicted, but cannot occur in practice.

Collectively, these findings establish that a simple model, relying exclusively on presence of m6A sequence motifs and exon-intron architecture, is able to recapitulate experimentally recorded m6A distributions along human and mouse genes.

### Exon-intron architecture dictates RNA stability via m6A

The strong dependency of m6A on exon-intron architecture prompted us to explore whether m6A was the missing link connecting exon-intron architecture and mRNA stability. We first sought to explore the extent to which m6Apred-2 was also predictive of mRNA stability. Remarkably, we found both in human and in mouse that predicted m6A levels correlated strongly with RNA stability, nearly as well as experimental counterparts. In human cells, there was a correlation of -0.5 between predicted m6A levels and experimentally measured half-lives in MCF7 cells (Schueler et al., 2014), close to the correlation observed between measured m6A levels (in A549 cells) and RNA stability (R_s_ = -0.57, **Fig. 4a**). Similarly, in mouse, we obtained a correlation of -0.47 between predicted m6A levels and experimentally measured half-lives in WT mESC cells (Ke et al., 2017), close to the agreement between stability and experimentally measured m6A levels (R_s_ = -0.50, **Fig. 4b)**. Critically, this correlation with half-life between either the predicted m6A level or the experimentally measured one are both abolished in METTL3 KO cells **(Fig. 4c)**.

**Figure 4.**
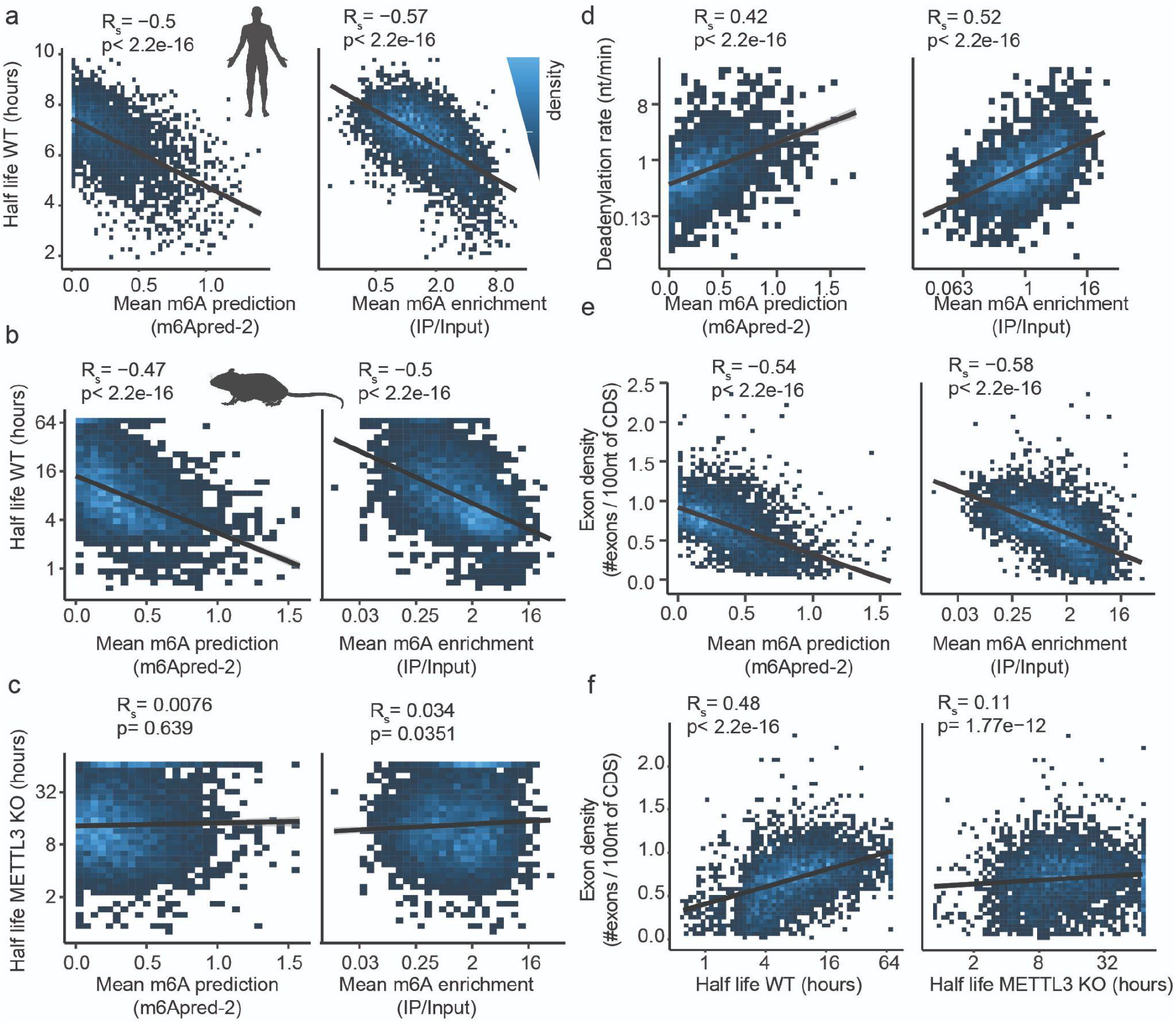
Exon-intron architecture drives mRNA stability through m6A. **a) (left)** Correlation between predicted m6A levels (m6Apred-2) and log-transformed mRNA stability rates in human cells. mRNA stability rates in MCF cells were obtained from (Schueler et al., 2014). **(right)** Correlation between measured m6A levels in A549 cells and mRNA stability. **b)** Plots as in (a), in mESC cells on the basis of predictions and measurements in mESC cells. **(c)** Corresponding plots of predicted and measured m6A levels in mESC cells against decay rates measured in METTL3 knock-out mESC cells. **d)** Correlation between predicted and measured m6A levels in mESC against deadenylation rates measured in mouse 3T3 cells. **e)** Correlation of exon density with predicted (left) and measured (right) m6A levels in mESC cells. **f)** Correlation of exon density with decay rate in WT and METTL3 KO mESCs. Mouse half-life values were capped at 72 h.

In a recent study, transcriptome-wide deadenylation rates were systematically inferred on the basis of measurements and modeling in mouse 3T3 cells (Eisen et al., 2020). Deadenylation rates were found to vary across 3 orders of magnitude across different genes, yet the basis for this variation remained unknown. Remarkably, we find that measured RNA deadenylation rates correlate well with predicted m6A levels (R_s_=0.42), close to levels achieved with measured ones (R_s_=0.52) **(Fig. 4d)**. These findings lend further support to the notion that m6A is a major factor contributing to mRNA deadenylation rates, consistent with findings that binding of m6A by YTH-domain harboring proteins results in deadenylase recruitment (Du et al., 2016).

In systematic studies exploring the determinants of mRNA stability, exon density within coding regions was found to be the primary correlate of mRNA stability (Agarwal and Kelley; Agarwal and Shendure, 2020; Sharova et al., 2009; Spies et al., 2013). We found that this metric correlates strongly with both measured (R_s_=-0.58) and predicted (R_s_=-0.54) m6A levels **(Fig. 4e)**, suggesting that exon density may simply be an indirect metric capturing m6A derived signal, given the depletion of m6A in exon-rich regions. To causally dissect this correlation, we explored the relationship between exon density and mRNA stability in WT and METTL3 KO mESCs. We could recapitulate a strong correlation between mRNA half-life and exon density (R_s_=0.48), which was strongly reduced in METTL3 KO mESC (R_s_=0.11) **(Fig. 4f)**. Thus, these results demonstrate that m6A is the underlying mechanism which connects exon-density with RNA half-life, and is as such by far the most dominant force known to shape RNA half-life.

Our findings establish strong quantitative associations between exon-intron architecture, mRNA methylation, mRNA deadenylation and mRNA degradation. The fact that these associations can be recapitulated by m6Apred-2 lends additional support to the validity of the model, and establishes a cardinal and predictable role for exon-intron architecture in dictating RNA methylation and mRNA degradation.

### Evolution of intron-exon architecture and m6A

Our results thus far suggest that our simple model, based on exon-intron architecture, has strong predictive power for human and mouse genes. We next sought to assess to what extent the rules giving rise to methylation were conserved across evolution. To this end, we analyzed m6A datasets in two additional species, zebrafish (Zhang et al., 2017) and Drosophila (Wang et al., 2021), in both cases on the basis of measurements in either WT or METTL3 depleted strains. Application of our model to zebrafish yielded a high agreement between experimentally measured m6A levels and predicted counterparts (R_s_=0.5, **Fig. 5a left**), which were dramatically reduced in zebrafish treated with METTL3-targeting morpholinos (R_s_=0.15, **Fig. 5a right**). These findings suggest that the mechanisms underlying m6A deposition are largely conserved between zebrafish and mammalian species. In stark contrast, in Drosophila we failed to obtain a correlation between measured m6A profiles and ones predicted based on our model (R_s_=0.007) **(Fig. 5b)**, suggesting that the underlying mechanism may have changed. If m6A distribution changed, this would predict that also the determinants of mRNA stability may have changed. As indicated above, mRNA stability in human and mouse is well correlated with both predicted and measured m6A levels. In contrast, in Drosophila we fail to see any association between experimentally measured mRNA half-lives in Drosophila embryos (Burow et al., 2015) and predicted m6A levels (R_s_=-0.04, p<1e-3, not shown). In these analyses we do observe a weak, yet significant, correlation with experimentally measured m6A levels (R_s_=0.21, p<2.2e-16, not shown), but this effect is maintained at identical magnitude in m6A measurements derived from METTL3 depleted flies (R_s_=0.23, p<2.2e-16, not shown), suggesting that this relationship is not m6A-dependent.

**Figure 5.**
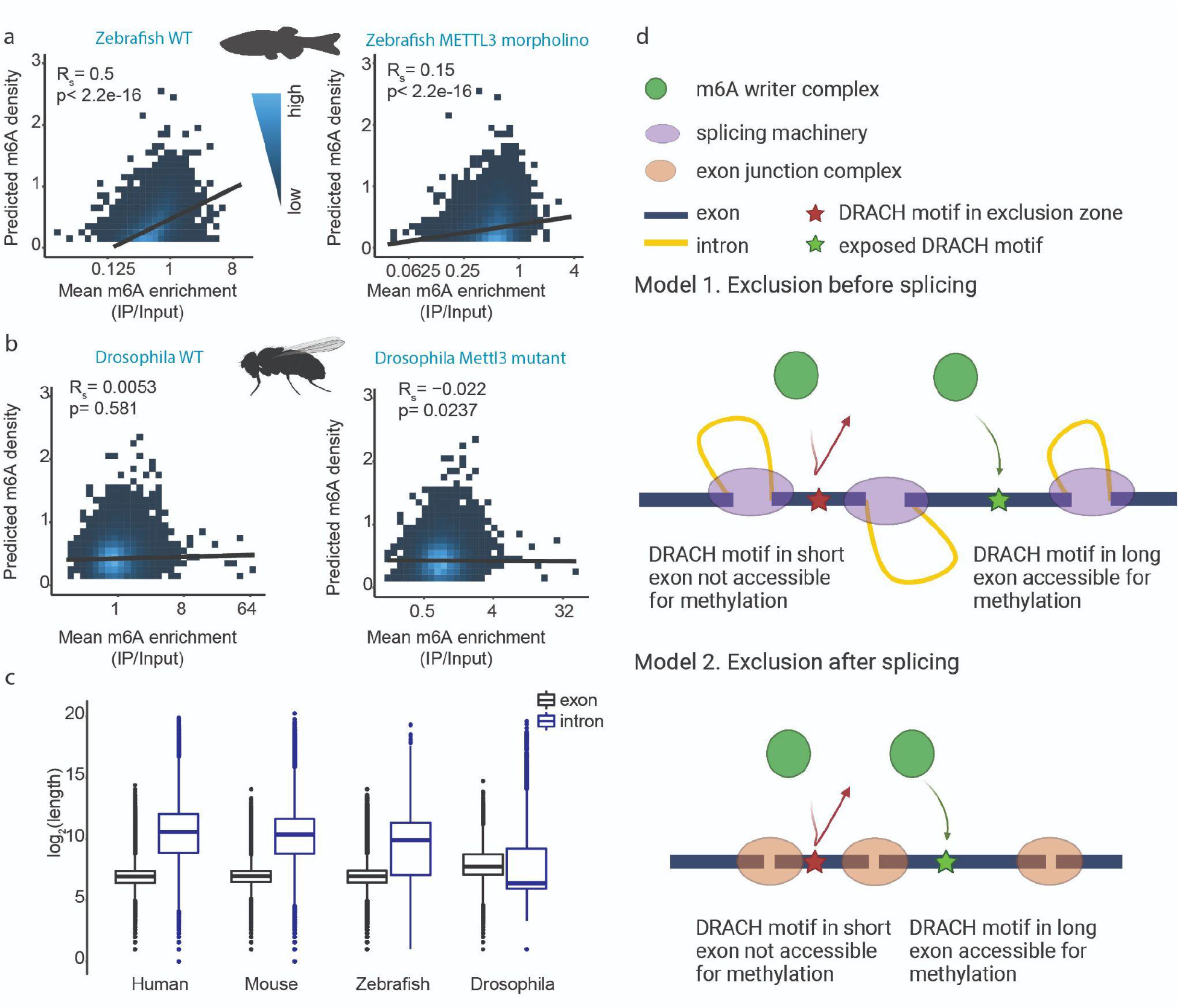
Evolution of exon-intron architecture is associated with evolution of m6A selectivity. **a)** Correlation between predicted m6A levels and m6A measurements in WT zebrafish embryos 28h post-fertilization (left) or METTL3-morpholino treated counterparts (right), obtained from (Zhang et al., 2017). **b)** Correlation between predicted m6A levels and m6A measurements in WT adult Drosophila (left) and corresponding METTL3 mutants (right), obtained from (Wang et al., 2021). **c)** Log-transformed lengths of all internal exons and introns in human, mouse, zebrafish and Drosophila. **d)** Possible models explaining the exclusion of methylation marks near splice junctions. Upper panel: exclusion of the m6A before or during splicing by the splicing machinery. Lower panel: exclusion of the m6A machinery after splicing by the exon junction complex (EJC) and/or EJC associated proteins.

These findings are of particular interest, given considerable changes in exon-intron architecture between human, mouse and zebrafish in comparison to flies. The former three have short internal exons spanned by long introns of variable length, whereas the latter has longer and more variable exons, but considerably shorter introns **(Fig. 5c)**. These differences in length are thought to reflect a difference in the mechanism via which introns are recognized and spliced by the splicing machinery, with ‘exon-definition’ being more prevalent in human, mouse and zebrafish with respect to ‘intron-definition’ in Drosophila (see **Discussion**). Our findings thus could potentially hint that evolution of m6A and mRNA stability profiles in higher eukaryotes may have been driven by evolution of exon-intron architecture and of the mode of intron recognition by the splicing machinery (see **Discussion**).

## Discussion

Here we unravel the mechanism by which m6A deposition is simultaneously dictated by sequence motifs and exon-intron architecture, and in doing so identify the mechanism linking exon-intron architecture to mRNA stability. Our findings have critical bearings on the biogenesis, regulation, function and evolution of m6A, on our understanding of the determinants, regulation and evolution of mRNA stability, as well as on our perception of splicing and its connection to downstream processes.

A fundamental challenge over the past decade has been to unravel the code giving rise to m6A. The vast majority of RNA modifications studied to date are catalyzed in a highly selective manner, typically at a specific set of sites in tRNA and rRNA. It was hence widely assumed that such a selective code must also be present for m6A, and potentially be encoded either in sequence, or in structure, or in some complex combination thereof. Herein we challenge this view, and suggest that m6A radically differs from other RNA modifications. Our findings suggest that m6A is installed in a non-selective, exclusion-based manner: all sites harboring a methylation consensus motif are methylated *unless* they are in proximity to a splice junction. This effectively segregates mRNAs into m6A-excluded zones, encompassing the vast majority of internal exons that tend to be short and hence cannot allow methylation, and m6A-permissive zones, within long exons (either internal ones or last exons), in which methylation consensus sites are found at a sufficient distance from splice junctions to allow them to undergo methylation. Interestingly, such an exclusion based manner is to some extent analogous to DNA methylation, which is largely exclusion based, with sites being methylated by default, *unless* they are inaccessible to the methylation machinery, for example due to DNA occlusion by a transcription factor (Hsieh, 2000).

Our study has critical bearings on our understanding of the determinants controlling mRNA decay. In a recent study in which mRNA decay patterns were analyzed across dozens of humans and mouse cell lines, it was found that the relative decay rates between different genes were to a large extent maintained both across different cell lines and across species (Agarwal and Kelley). The primary feature with which these ‘fixed’ decay rates correlated was exon density, in line also with prior studies (Agarwal and Shendure, 2020; Clark et al., 2012; Sharova et al., 2009; Spies et al., 2013). The underlying mechanism, connecting (nuclear) splicing and (cytoplasmic) decay had been unknown. Here we reveal that the basis for this association is m6A - a covalent, co-transcriptionally introduced mark - whose deposition we uncover to rely on intron-exon architecture, and which mediates cytoplasmic degradation. Of note, the potential for a link between methylation and splicing has been explored by several studies, but all through an inverse lens, of whether methylation *impacts* splicing (Ke et al., 2017; Lence et al., 2016; Wei et al., 2021; Xiao et al., 2016); these studies typically found a limited impact. In contrast, our results suggest that the impact of splicing on methylation is profound; Indeed, we find that exon-intron architecture is the primary force sculpting region-specific methylation topographies. As a hard-wired mechanism reflecting exon-intron architecture, our findings also offer an attractive lens through which to re-examine additional contexts in which the cytoplasmic fate of an mRNA (e.g. localization, translation) were found to be associated with (nuclear) splicing (Buchman and Berg, 1988; Dwyer et al., 2021; Le Hir et al., 2001, 2003; Luo and Reed, 1999; Matsumoto et al., 1998; Proudfoot et al., 2002; Zuckerman et al., 2020). The mechanisms proposed for such observations included RNA binding proteins that can bind cotranscriptionally and shuttle from the nucleus to the cytoplasm and the deposition of the exon junction complex at exon-exon junctions, but these only partially account for the observations.

Our findings have widespread implications on the regulation of m6A and of mRNA stability. The majority of exons are constitutively spliced most of the time. As such, mechanisms tied to splicing are similarly expected to be constitutive, to a considerable extent. This inference is consistent both with the relative robustness of m6A profiles across different cell and tissue types (Liu et al., 2020b; Schwartz et al., 2014) as well as with recent findings that mRNA stability is also highly correlated across diverse cell types (Agarwal and Kelley). Nonetheless, exon-intron architecture can be modulated via different forms of alternative splicing or via alternative polyadenylation, which can alter the proximity of m6A consensus sites to splice junctions, thereby impacting methylation potential and, in turn, mRNA stability. Alternative polyadenylation events are particularly interesting to consider, as it has been observed in diverse contexts that longer isoforms are less stable. This reduced stability was attributed to abolishment of microRNA binding sites, though this abolishment was shown to only account for a minor portion of this destabilization (Mayr and Bartel, 2009; Sandberg et al., 2008). Our results offer a fresh interpretation for these observations, given that longer 3’ UTRs give rise to broad methylation-permissive regions (vertebrate 3’ UTR typically lack introns), which increase the methylation load of a gene and thereby reduce stability.

Our findings are also of bearings on m6A evolution, and how it may be tied to evolution of splicing. Human, mouse and zebrafish all harbor a similar exon-intron architecture, comprising short internal exons relatively narrowly distributed around an average length of ∼130 bp and much longer introns of variable lengths (Schwartz et al., 2008). In these species, it is thought that the primary unit that is recognized by the splicing machinery is often the exon (‘exon-definition’) (Barutcu et al., 2022; Tammer et al., 2022), and that the splicing factors hence initially assemble on both sides of the exon, constraining its length. In contrast, in Drosophila the splicing machinery is thought to primarily assemble along both ends of the intron (‘intron definition’) (Keren et al., 2010), and consistently exons are more variable in length, and intron length is more constrained. Our model has high predictive power in the three species subscribing to an exon-recognition regime, but lacks predictive power in Drosophila, subscribing to an intron-definition regime. Our model also has strong predictive power of mRNA stability in human and mouse, subscribing to an exon-definition regime, but lacks such power in Drosophila. It is tempting to speculate that the differences in the rules governing m6A formation and mRNA stability are linked to this fundamental divide in exon-intron architecture and to the different mechanisms via which introns in these different species are recognized and spliced.

Our findings suggest that rather than being present on only a handful of sites per gene, m6A is likely installed at hundreds of thousands of sites transcriptome-wide, with an average of ∼13 m6A sites per gene. Such numbers are not consistent with original estimates of 2-4 sites per gene (Perry and Kelley, 1974) - however, these estimates had relied on an assumption that m6A sites are stoichiometric, which is unlikely to be the case for the vast majority of sites. These numbers are also dramatically higher than the ∼5,000-50,000 ‘peaks’ which were typically reported in individual m6A-seq experiments (Dominissini et al., 2012; Ke et al., 2015; Körtel et al., 2021; Meyer et al., 2012) - the discrepancy, in this case, is likely due both to the difficulty of ‘peak calling’ and due to the fact that, as we now show, a single peak is typically the product of multiple m6A sites. The numbers of methylation sites that we report are consistent with inferences previously made by us on the basis of antibody-independent measurements in human and yeast that led us to conclude that original estimates of m6A sites had been substantially underestimated (Garcia-Campos et al., 2019). These numbers are also in general agreement with recent results based on monitoring of m6A in single cells (Tegowski et al., 2022). These numbers have important bearings on the utility of probing the function of individual m6A sites. Given the wide spread of methylation, the multiple methylation sites contributing to individual peaks, and the low stoichiometries of individual sites, the functional impact of any individual methylation site may often be marginal. The relevant resolution with which to consider m6A, and at which m6A sites need be perturbed for functional studies, may not be single-base, but instead of entire genes.

The precise mechanism by which splicing prevents methylation from occurring at junction-proximal regions remains to be defined. This question is intricately related to the relative timing of m6A deposition with respect to mRNA splicing, which is not fully understood. On the one hand, m6A appears to be present at near steady-state levels already on chromatin-associated RNA (Ke et al., 2017), is present on nascent RNA (Xu et al., 2022), and was found by one study to be widespread on introns (Louloupi et al., 2018), suggesting that it is deposited prior to splicing. Yet, most studies have found the modification to be strongly depleted from introns (Chen-Kiang et al., 1979; Ke et al., 2017; Wei et al., 2021), suggesting that it may be installed following splicing. If m6A deposition precedes splicing, then it is conceivable that the splicing machinery, a multi-megadalton complex comprising dozens of proteins (Brody and Abelson, 1985; Will and Lührmann, 1997), physically occludes the junction-proximal sequence, and prevents access of the methylation complex, which is itself over a megadalton in size (Bokar et al., 1997) **(Fig. 5d, top)**. Alternatively, if m6A is deposited only following splicing, its depletion from junction-proximal regions could reflect potential occlusion via exon junction complex and associating RNA binding proteins (Le Hir et al., 2016; Merz et al., 2007; Singh et al., 2012) **(Fig. 5d, bottom)**.

It is also interesting to speculate about the ‘logic’ of coupling mRNA degradation to absence of splice junctions. One possibility is that such coupling came about in order to inhibit gene expression from retrotransposons. Such elements are pervasive in the human genome, they lack introns, and their expression has the potential to create havoc. Retrotransposons are epigenetically silenced, at the transcriptional level, via 5mC on DNA - indeed, 5mC is speculated to have evolved as a mechanism for their silencing (Deniz et al., 2019; Yoder et al., 1997). m6A, which - similar to m5C - is installed by default and imposes silencing but at the RNA level, may serve as a post-transcriptional counterpart. This idea resonates with recent literature implicating m6A in the silencing of transposable elements (Chelmicki et al., 2020; Chen et al., 2021; Liu et al., 2020a, 2021; Xu et al., 2021).

Currently, our model exhibits a correlation of up to R=∼0.7 with experimentally measured methylation levels. While substantial, this only accounts for ∼50% of the variability in experimentally measured m6A levels. There are likely several factors underlying this partial correlation. First, the experimental measurements - relying on an anti-m6A antibody - are an imperfect proxy of actual methylation levels, given antibody promiscuity and bias (McIntyre et al.; Schwartz et al., 2013). Second, the genomic annotations - on which the model heavily relies - are an additional source of error. 5’ and 3’ UTRs are particularly variable, and often there can be substantial discrepancies between annotations of transcript starts and ends versus the actual starts and ends in individual cell lines. Such discrepancies can lead to substantial differences between predicted and actual methylation levels, and may in part underlie the fact that the agreement between predictions of our model and experimentally measured data in the CDS and beginning of 3’ UTR is substantially higher than across the distal end of the 3’ UTR. Third, a strength - but also weakness - of our model is its simplicity: it relies on binary decisions regarding the eligibility of motifs and junction-distances. We anticipate that more sophisticated approaches, including more quantitative modeling of the motif and the distances, and potentially also factoring in additional parameters such as secondary structure (Garcia-Campos et al., 2019; Meiser et al., 2020; Schwartz et al., 2013), will further boost the agreement with measured m6A levels.

Collectively, our findings provide a unifying model, quantitatively and causally connecting exon-intron architecture, m6A, and RNA stability. We anticipate that this study will open up future explorations into the mechanism that prevents m6A formation in the vicinity of a splice junction, into the logic of tying mRNA stability to exon-intron architecture, and into physiological and pathological contexts in which regulation of exon-intron architecture serves as a mechanism for regulating mRNA stability by regulation of the deposition of m6A.

## Materials and methods

### Cloning

SLC25A3 WT and mutant gene fragments (Table S1) were ordered from Twist Biosciences and cloned with restriction-ligation based approach to the 3’ end of GFP using BcuI (Thermo Fisher Scientific) and BstBI (NEB). An oligo library pool was synthesized by Twist Biosciences and cloned as detailed in (Uzonyi et al., 2022). The full list of sequences in the pool can be found in Table S2.

For the wild type sequences, 200 human and 200 mouse genes were selected based on an annotated methylation site in several miCLIP data sets, aiming to keep a variability of different sequence motifs, as well as different origins of the sites, such as internal exons, end of CDS and 3’ UTR. A 50 bp region on both sides of the methylated adenosine was selected based on the hg19 and mm9 genome annotations, leading to a total of 400 sequences with 101 nt length. For the mut-main set, the adenosine at position 51 was mutated to a T. For the mut-secondary set, all As in a DRACH motif, except for position 51 were mutated to a T. For the mut-all set, all As in DRACH motifs were mutated to a T.

The ‘Running mutation’ series is based on 10 human mut-secondary sequences. In this series, 1, 3 or 5 consecutive bases were mutated to their complement, base-by-base, from the first to the last of the 101 nucleotides. The ‘Sequence mutation’ series is based on 4 human mut-secondary sequences. In this set, the extended, 9-mer consensus motif in the middle of the sequence was permuted to all possible combinations of NTNRACNGN, with the methylated A in the middle. The ‘Running methylation’ set is based on ten human mut-secondary sequences. In this set, the extended 9-mer motif was moved from the beginning to the end of the 101 nt sequence, base-by-base. The ‘Motif number’ seq is based on 18 different 9-mer m6A consensus motifs: ATAAACAGT, ATAAACCGT, ATAAACTGT, ATAGACAGT, ATAGACCGT, ATAGACTGT, ATGAACAGT, ATGAACCGT, ATGAACTGT, ATGGACAGT, ATGGACCGT, ATGGACTGT, ATTAACAGT, ATTAACCGT, ATTAACTGT, ATTGACAGT, ATTGACCGT and ATTGACTGT. Each motif was placed in the 101 nt sequence in its wild type form n times (0-8) and in its point mutant (T instead of A in the middle) form 8-n times, at random locations. The remaining bases were filled up with random nucleotides keeping a 50% GC content.

Each sequence was planned with a unique 10 nt barcode. Barcodes were designed to be devoid of ‘AC’ dinucleotides, to avoid any chance of undergoing methylation, and of purine triplets, as purine rich regions may be promiscuously immunoprecipitated by the anti-m6A antibody (Schwartz et al., 2013). Furthermore, barcodes and the planned sequences were designed to be devoid of canonical poly-adenylation signals and the restriction sites used in the cloning strategy.

DNA sequences encoding the mice Nol12, Coa3 and Calm3 genes were ordered from Twist Bioscience. These initial sequences included the full respective ORFs with either 1) the last intron of each gene (three introns for Calm3), 2) point mutations of adenosines suspected to be m6A-modified or, 3) no introns. These sequences were PCR-amplified and introduced between HindIII and NotI sites in the pcDNA5/FRT/TO plasmid (Thermofisher). For subsequent mutagenesis or modifications, the desired changes were introduced using restriction-free cloning. In brief, the desired changes were introduced using primers with partial complementarity to the original construct being modified and PCRs producing two parts of the new construct with complementary overhangs were conducted. After isolation of the correct products from agarose gels, both PCR products were combined and amplified using terminal primers. The final products were isolated from gels, digested with HindIII and NotI and ligated into a linearized pcDNA5/FRT/TO plasmid (Thermofisher). All PCRs were performed using Kapa HiFi DNA polymerase (Roche). Final constructs were partially sequenced using Sanger sequencing. All final sequences are described in Table S1.

### Cell culture and transfections

V6.5 mESC line was kept in DMEM (Gibco) supplemented with 1% Penicillin-Streptomycin, 1 mM L-glutamine, 1% non-essential amino acids, 20% high-grade fetal bovine serum (Biological Industries), beta-mercaptoethanol and 10 µg recombinant leukemia inhibiting factor. Cells were kept on tissue culture plates covered by gelatin (0.2%), in co-culture with in-house generated, radiation-inactivated mouse embryonic fibroblasts. Cells were seeded on gelatin-covered 6-well plates (without MEF feeder cells) for transfection. GFP-SLC25A3 plasmid was transfected with the TransIT-X2 reagent (Mirus) according to the manufacturer’s instructions. Cells were collected 24 hours post-transfection.

HEK293T cells were cultured in DMEM (GIBCO) supplemented with 10% FBS and 1% Penicillin and Streptomycin. For the library pool transfection, 0.5 million HEK293T cells were seeded in each well of a 6-well plate. 24 hours later cells were transfected with 2 μg plasmid DNA and 8 ul home-made PEI reagent. Cells were harvested 24 hours post transfection. Each 3 wells of the 6-well plate were merged to form two replicates.

MCF7 cells (ATCC) were maintained in DMEM (GIBCO) supplemented with 10% FBS and 1% Penicillin and Streptomycin. Transfections were done with JetPrime (Polyplus) reagent, according to the supplier’s instructions. Typically, the cells were seeded in 6-cm dishes (Corning) and on the next day transfected with 1-2µg of plasmids. The cells were harvested after additional 24 hours by scraping and subjected to RNA extraction.

### RNA extraction, m6A IP and library preparation

RNA from mESC and HEK293T cells was extracted with NucleoZOL (Macherey-Nagel); from MCF7 cells BIO TRI RNA reagent (Bio-lab) was used. RNA was poly-A selected with oligo dT-beads (Dynabeads mRNA DIRECT Kit life tech). For all single-plasmid transfections, sample pooling, m6A immunoprecipitation and NGS library preparation was prepared according to the step-by-step m6A-seq2 protocol described in (Dierks et al., 2021). m6A IP and amplicon library preparation of the oligo library pool was performed on the basis of the same protocol with the following modifications: the 3’ adapter ligation and pooling steps were omitted, the 5’ adapter ligation step was omitted, reverse transcription was performed with a sequence specific RT primer (Table S1), library amplification was performed with sequence specific PCR primers containing Illumina sequencing adapters (Table S1). All NGS libraries were sequenced on the Illumina NovaSeq 6000 platform.

### Data analysis

Oligo pool data was analyzed with a custom R script. Enrichment scores were calculated as the ratio between the number of reads originating from a single sequence upon m6A-IP in comparison to the input and normalized to the median score of all human, mouse and synthetic constructs with no m6A consensus motif in the library.

Transcriptome wide m6A-seq2 datasets were mapped to mm9 (mESC) or hg19 (MCF7) genome assembly and the transfected plasmid with STAR/2.7.9a (Dobin et al., 2013). Per-base read coverages were obtained with txtools (https://github.com/AngelCampos/txtools), and normalized to the total number of reads in the pool. Raw input and IP read counts were first normalized by the sum of reads across the entire corresponding pool of samples subjected to m6A-seq simultaneously (i.e. across all input samples or all IP samples, respectively). m6A enrichment levels were calculated as the fold-change in normalized coverage at the modified position between IP and input.

For constructs with large insertions and deletions (in **Fig. 2e-d**), we sought to avoid contribution of signal originating from the insert (or from upstream intronic regions). In these analyses we therefore constrained the analysis of input and IP reads only to ones overlapping the sites and beginning at most n bases upstream of the methylated position.

In the case of **Fig. 2e**, n=10 was employed, given the presence of a DRACH motif 10 nt upstream of the shortest construct (in which the targeted m6A site was 29 nt away from the splice junction). To quantify the constructs with intron retention **(Fig. 2f)**, n=40 was employed; In this case a longer distance could be used, allowing integration of more signal, without reaching the DRACH site.

### Datasets analyzed in this study

Measurements of m6A in WT and WTAP-depleted A549 cells were obtained from (Schwartz et al., 2014) (GSE54365). Measurements of m6A in WT and METTL3 KO mESC cells were obtained from (Garcia-Campos et al., 2019), (GSE122961, replicate 1). Measurements of m6A in WT and METTL3-morpholino treated zebrafish embryos (28 hours post-fertilization) were obtained from (Zhang et al., 2017) (GSE89655). Measurements of m6A in WT adult Drosophila and corresponding METTL3 mutants were obtained from (Wang et al., 2021) (GSE155662). mRNA stability in WT human MCF7 cells were obtained from (Schueler et al., 2014) (GSE49831). mRNA stability in WT and METTL3 KO mESC cells were obtained from (Ke et al., 2017). mRNA stability in Drosophila embryos were obtained from (Burow et al., 2015). Deadenylation rates were obtained from (Eisen et al., 2020).

### Implementation of m6Apred-1 and m6Apred-2 models

Gene models for roughly 20,000 ‘canonical’ human and mouse genes were downloaded from the ‘UCSC Known Genes’ annotation table in the UCSC genome browser. For each gene we identified all ‘eligible’ DRACH motifs, and furthermore recorded their distance from the nearest exon-intron junction. In m6Apred-1, each of the motifs was considered methylated. In m6Apred-2, an eligible DRACH motif was only considered methylated if its distance to the nearest splice junction exceeded 100 nt. To mimic the regional enrichment in m6A-seq, every site predicted to undergo methylation was modeled as a gaussian over a 200 bp region centered at the methylated site. The values along this gaussian were derived using the density function of a gaussian distribution (mean=0, sd=4), which were calculated for 100 values distributed at fixed intervals between 0 and 9 using the dnorm() function in R, which were min-max normalized, to distribute between 0 and 1. The final predicted enrichment value at each position along the gene was defined as the sum of all signal (stemming from zero, one or potentially multiple gaussians) overlapping this position.

For the analyses summarizing gene-level predicted or measured methylation levels, we calculated the mean signal over gene bodies. Gene bodies were defined as the entirety of the gene excluding TSS-proximal regions (first 200 bp of the gene) and distal 3’ UTR regions (greater than 400 bp from the stop codon). These regions were excluded to avoid contribution from signal originating from m6Am at the beginning of genes (Akichika et al., 2019; Keith et al., 1978; Mauer et al., 2016; Schwartz et al., 2014) and limitation in available annotations of 3’ UTRs (see Results). The gene-level quantification of experimentally-measured m6A levels were calculated as the overall number of reads per gene-body in the IP experiment divided by the corresponding number in the input experiment. This metric draws on our previously developed m6A-gene index (GI) (Dierks et al., 2021).

### Analysis of experimental m6A-seq datasets and correlations against predicted values

m6A-seq measurements (IP and input) were downloaded from the sources above, and aligned against the hg19, mm9, danRer11 and dm3 genomes for human, mouse, zebrafish and Drosophila, respectively, using STAR (Dobin et al., 2013). An in-house script was employed to map the genomic coordinates to transcriptomic annotations. For these annotations, we used canonical ‘UCSC Known Genes’ for human and mouse, Ensemble for zebrafish, and RefSeq for Drosophila. Only reads fully matching a transcript structure were retained. For all data generated using m6A-seq and m6A-seq2 which provides paired-end data (note: this includes all employed data with the exception of the zebrafish and Drosophila datasets), reads were computationally extended in transcriptome space from the beginning of the first read to the end of its mate, and coverage in transcriptome-space was calculated for each nucleotide across all transcripts. For each base along each transcript, an enrichment score was calculated, defined as the IP/Input. For assessing agreement between predicted and experimentally measured profiles, we only included protein-coding genes with an average coverage per base exceeding 5 reads in the Input sample, to ensure adequate expression for estimating enrichment levels. These filters typically allowed quantifications of ∼5000-∼10000 genes.

## Supporting information

Table of MPRE sequence information

Table of primer and oligo sequences

## Supplementary Figures

**Figure S1.**
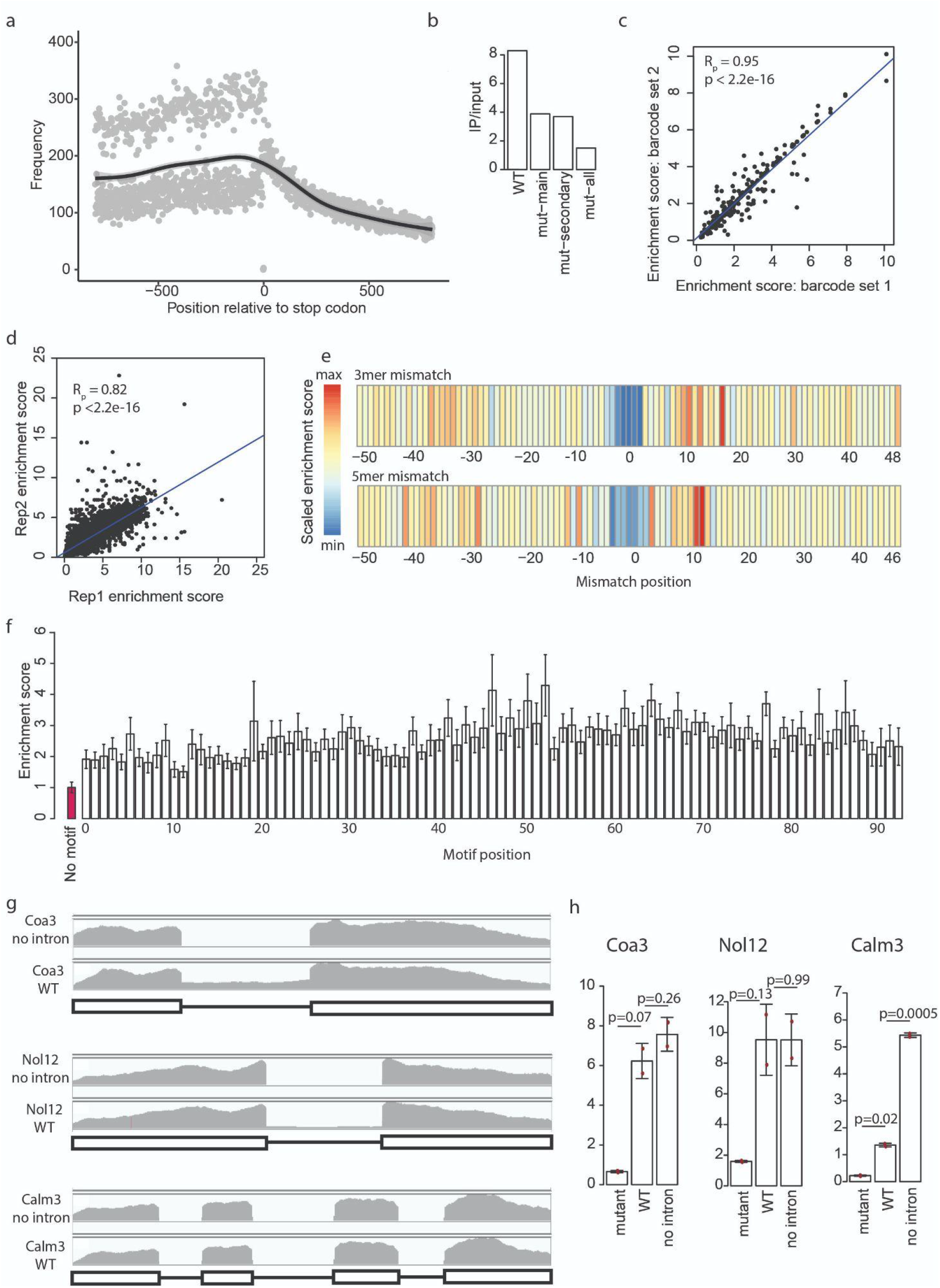
**a)** Metagene plot of the distribution of the 7 most common DRACH motifs (GGACC, AGACA, TGACT, AGACT, GAACT, GGACA, GGACT) in the human genome. Black line shows Loess fitted average. **b)** IP/input ratio from panel Fig. 1a, at the position of the main consensus motif, based on a single measurement. **c)** Correlation of the enrichment score of 150 sequences planned with two different barcodes. Enrichment score was calculated by dividing the IP reads ratio mapped to a barcode by the input reads ration mapped to a barcode (IP/input ratio), and normalized to the median IP/input ratio of all human, mouse and synthetic sequences with no DRACH motives present in the pool. Pearson correlation coefficient and p-value are marked on the plot. **d)** Correlation of biological replicates of the MPRA assay. Pearson correlation coefficient and p-value are marked. **e)** Enrichment scores in the running 3mer and 5mer nucleotide point mutation, according to Fig. 1e. The mismatch position marks the first mutated base. **f)** Enrichment scores of the moving motif series. 9-mer m6A motif was moved base-by-base from the beginning to the end of the 101 nt long sequence. Bar plot depicts the mean and standard error of the mean. The value for the negative control sequences with no DRACH motif is marked in red. **g)** IGV coverage profiles of the WT (intron harboring) and intronless sequences of the Coa3 (upper panel), Nol12 (middle panel) and Calm3 (lower panel) constructs, showing splicing of the WT sequences. Black boxes mark exons, horizontal lines mark introns. **h)** IP/input ratios at the position of the main DRACH motif of the Coa3, Nol12 and Calm3 constructs related to Fig. 1h-j. Red dots mark individual measurements, the height of theh-j bars mark the average and the whiskers represent the standard deviation of the mean. P-values for two-tailed Student’s t-test are shown.

**Figure S2.**
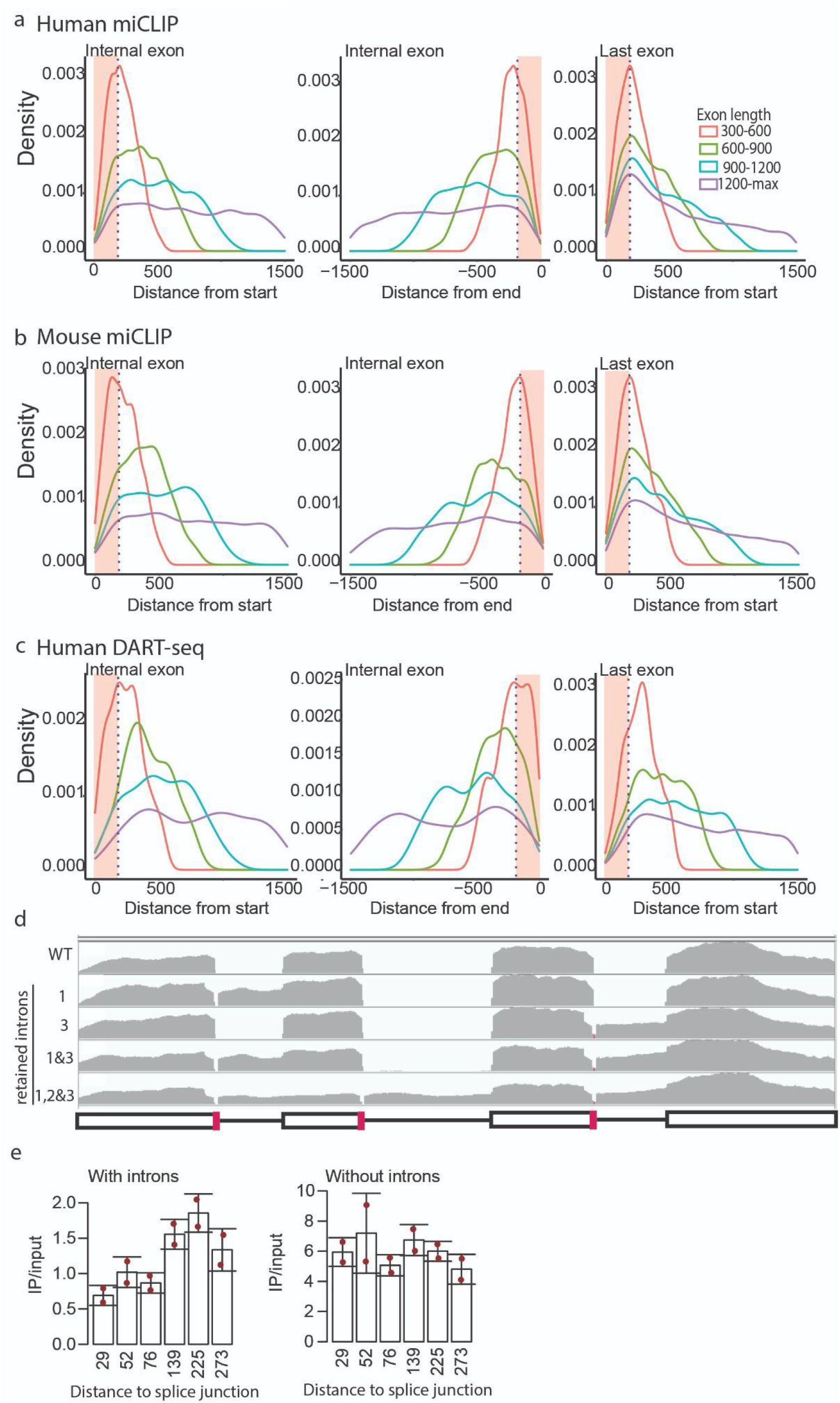
**a)** Histogram of detected methylation sites in human miCLIP data based on exon length, according to Fig. 2d. While Fig. 2d only shows exons over 600 bp and the first/last 500 bp of exons, this figure shows all exons over 300 bp in a distance of up to 1500 bp from the splice junction. Left panel shows distance from the start of internal exons, middle panel: distance from the end of internal exons, right panel: distance from the start of last exons. **b)** As in b), on the basis of mouse miCLIP data. **c)** As in b), on the basis of human bulk DART-seq data. **d)** IGV coverage profiles of WT (intron harboring) Calm3 construct, and corresponding constructs exhibiting retention of intron 1, 3, 1&3 or 1,2,&3, due to 5 nt deletions at the splice junction. Black boxes mark exons, horizontal lines mark introns, red marks the 5 nt deletion leading to intron retention. **e)** Related to Fig. 2e, bar plots comparing the methylation levels of the 3’ UTR DRACH site of the WT Calm3 construct with or without introns, and corresponding 5’ read starts until the nearest m6A site in the shortest construct (10 nt) were summed. The quantification rationale is further explained in the methods section. Red dots mark individual measurements, the height of the bars mark the average and the whiskers represent the standard deviation of the mean.

**Figure S3.**
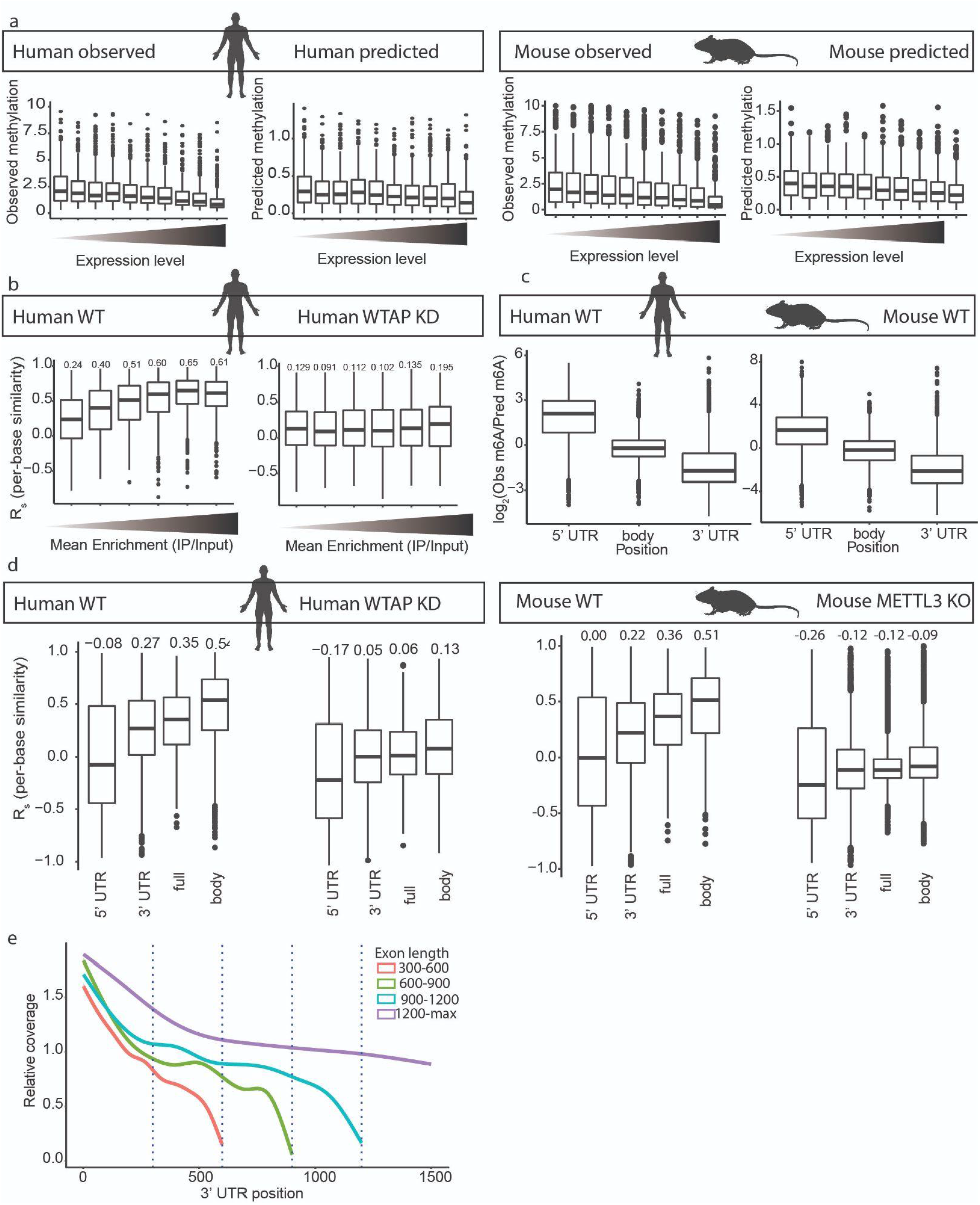
**a)** Box plots showing the relation between expression level and experimentally observed methylation in human (left panel), m6Apred-2 predicted methylation in human (middle-left), experimentally observed methylation in mice (middle-right) and m6Apred-2 predicted methylation in mice (right). Box plots correspond to the median, Q1 and Q3, whiskers mark Q1-1.5 IQR and Q3+1.5 IQR. Genes are divided into ten equally sized bins according to their expression levels. **b)** Human WT and WTAP KD counterpart of the data in Fig. 3f. **c)** Box plots of observed versus m6Apred-2 predicted m6A in human (left) and mouse (right) in different genetic regions; 5’ UTR, gene body and 3’ UTR. **d)** Box plots of the m6Apred-2 model performance in human (left) and mouse (right) in different genetic regions. Y-axis shows Spearman correlation coefficient of per-base similarity. Median correlation coefficients of each box are written over each box. **e)** Metagene coverage plots along the last annotated exon within genes (harboring the 3’ UTR), binned based on annotated length of the last exon. Substantial drops in coverage are observed long before the annotated ends, suggesting that the actual 3’ UTR is typically shorter than its annotated version.

**Figure S4.**
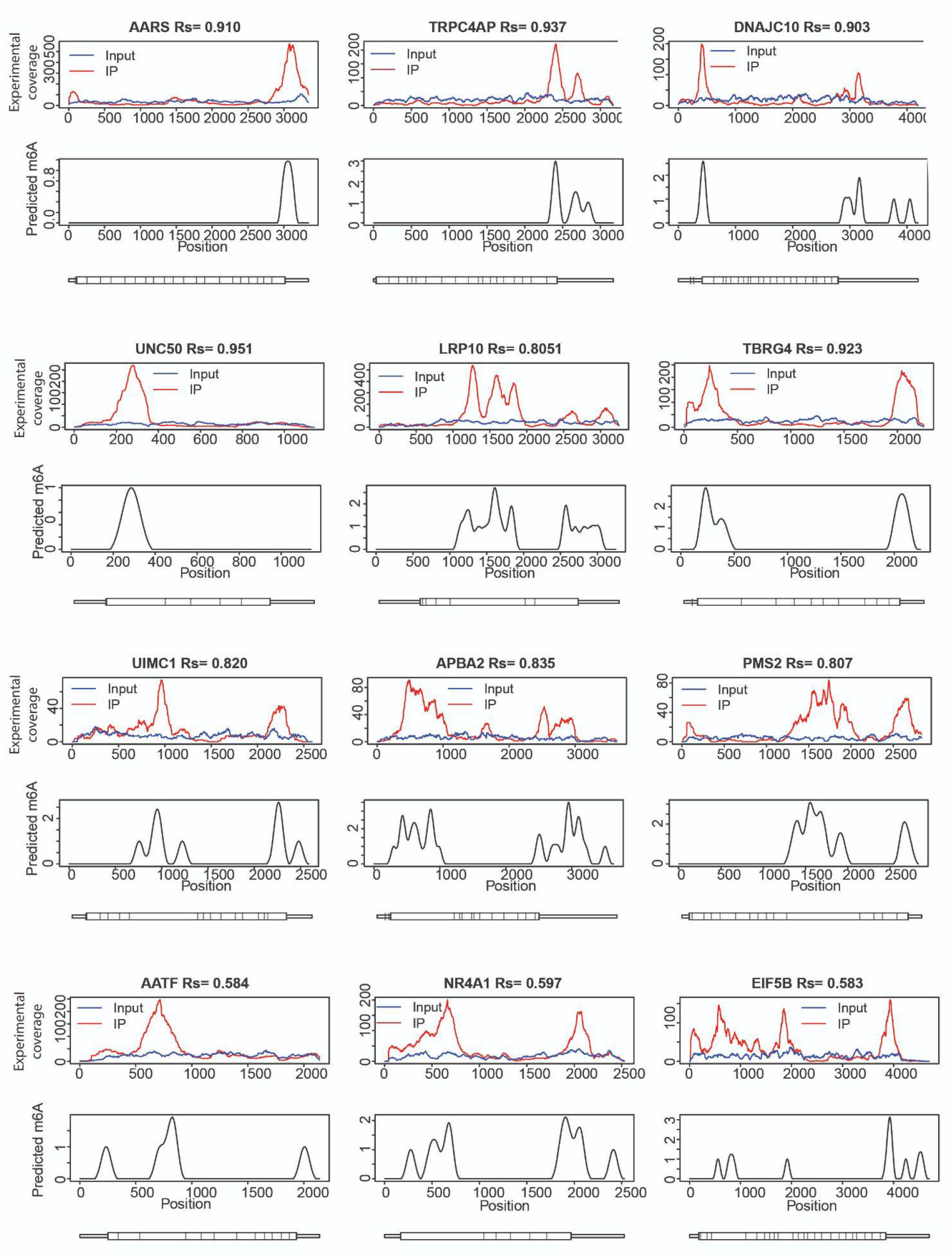
a) Predicted and measured m6A levels across 12 additional human genes, as in Fig. 3e.

